# Evaluation of the OsTIR1 and AtAFB2 AID systems for chromatin protein degradation in mammalian cells

**DOI:** 10.1101/2021.03.15.435352

**Authors:** Anastasia Yunusova, Alexander Smirnov, Tatiana Shnaider, Svetlana Afonnikova, Nariman Battulin

## Abstract

Auxin-inducible degron (AID) system is a promising tool for dynamic protein degradation. In mammalian cells, this approach has become indispensable to study fundamental molecular functions, such as replication, chromatin dynamics or transcription, that are otherwise difficult to dissect. We present evaluation of the two prominent AID systems based on OsTIR1 and AtAFB2 auxin receptor F-box proteins (AFBs). We analyzed degradation dynamics of cohesin/condensin complexes subunits in mouse embryonic stem cells (mRad21, mSMC2, mCapH, mCapH2) and human haploid HAP1 line (hRad21, hSMC2). Double antibiotic selection helped to achieve high homozygous AID targeting efficiency for all genes, ranging from 11 to 77%. We found that the main challenge for successful protein degradation is obtaining cell clones with high and stable AFB expression levels due to mosaic expression of AFBs, which also tends to decline with passages in the absence of constant puromycin selection, even at the AAVS1 safe-harbor locus. Comparing two AFBs, we found that OsTIR1 system showed weak dynamics of protein degradation. At the same time, AtAFB2 approach was very efficient even in random integration. Other factors such as degradation dynamics and low basal depletion were also in favor of AtAFB2 system. Our main conclusion is that repeated addition of puromycin to culture medium prevents AtAFB2 silencing and restores auxin sensitivity, facilitating robust protein degradation. We hope that our report will be useful for researchers that plan to establish AID method in their lab.

## INTRODUCTION

Loss-of-function experiments reveal important information about molecular pathways and gene functions. For most critical genes, whose knockout is lethal for cells, temporal knockdown methods, such as RNA interference, are employed, but could be ineffective. Auxin-inducible degron (AID) system has provided research community with a unique tool for rapid and complete degradation of a protein of interest (POI), controlled by addition of auxin molecule [1]. AID system has been successfully applied in many organisms ranging from yeast to transgenic mice [2]. Adaptation of the AID system in popular cell cultures, such as HeLa, HCT116, mouse and human stem cells, neurons [3–8], was extremely fruitful and allowed to study complex molecular pathways for virtually unlimited applications.

To perform AID experiments, cells should be modified with specific genetic constructs. First, a short peptide AID degron domain is fused via endogenous modification to a protein of interest (POI) at the N-or C-terminus. In addition, an auxin receptor F-box protein (AFB) is expressed to form an ubiquitin E3 ligase complex with other cellular components. AFB recognizes AID domain of the chimeric POI in the presence of the auxin molecule (indole-3-acetic acid (IAA) or other analogs), which leads to polyubiquitination and fast proteasomal degradation of the POI, usually over the course of just a few hours.

Researchers have employed several AID systems using different AFBs, of which *Oryza sativa* TIR1 (OsTIR1) [9] has become the most popular choice; and *Arabidopsis thaliana* AFB2 (AtAFB2), that has come to prominence in recent years [10]. OsTIR1 facilitates degradation of POI tagged with miniAID domain (amino acids 68–132 of *Arabidopsis thaliana* IAA17 protein), while AtAFB2 interacts with miniIAA7 domain (amino acids 37–104 of the same protein), leading to similar results. Experimental data show that AtAFB2 approach could be preferential due to faster degradation dynamics and lack of basal degradation (depletion of the POI in the absence of auxin) [10].

During last years, researchers have come up with several strategies to generate stable transgenic cell lines expressing AID system. But even though many great protocols exist [9,11,12], the method still contains pitfalls that could obstruct new users. For instance, selection of the proper transgenic clone with high POI degradation efficiency could be unmanageable due to AFB silencing. We decided to share our experience of using two popular auxin degron systems to help readers to avoid some common difficulties.

In our experiments, we used AID system to study effects of condensin/cohesin complexes depletion on nuclear architecture in two popular cell types: mouse embryonic stem cells (mESCs) and human haploid HAP1 cell line. For this purpose, we tagged Rad21, SMC2, CAPH and CAPH2 genes with miniAID tag (OsTIR1 system) [3] or miniIAA7 tag (AtAFB2 system) (Li et al., 2019), transfected cells with corresponding AFBs, and analyzed protein degradation dynamics, using microscopy, FACS and Western blotting.

## RESULTS

### Gene locus targeting with AID cassette in mES and HAP1 cells

We tagged endogenous target loci with AID degron tag fused with Clover/eGFP by CRISPR/Cas9-mediated HDR (Natsume et al., 2016; Li et al., 2019). In total, we modified 4 genes (Rad21, SMC2, CAP-H, CAP-H2) in mES cells; and 2 genes (Rad21, SMC2) in HAP1 cells (Table 1). We used a single gRNA targeting a region around the STOP codon to introduce AID degron and selection cassette at the 3’-ends of the genes (Fig. 1). Homology-directed integration of the AID cassette was facilitated by 600-1300 bp homology arms (electronic supplementary material Table S2). In our study, we used a minimal required AID degron (either mAID or miniIAA7) linked to Clover/eGFP (Fig. 1). Donor vectors also contained one of the two selection markers (Neomycin^R^/Hygromycin^R^) [3], which allowed to achieve efficient targeting for all genes with double selection.

**Table 1.** Gene targeting efficiencies in mESCs and HAP1 clones on the example of AtAFB2 system.

|  | **Total colonies**<br>**genotyped** | **Homozygous targeting** | **Mutations (deletions/**<br>**insertions)** |
| --- | --- | --- | --- |
| ES-Rad21 | 15 | 5 (33.3%) | 3 (20%) |
| ES-SMC2 | 15 | 6 (40%) | 3 (20%) |
| ES-CapH | 24 | 6 (25%) | 4 (16.7%) |
| ES-CapH2 | 19 | 5 (26%) | - |
| Hap1-Rad21 | 37 | 4 (11%) | 1 (3%) |
| Hap1-Smc2 | 13 | 10 (77%) | 2 (15%) |

**Figure 1.**
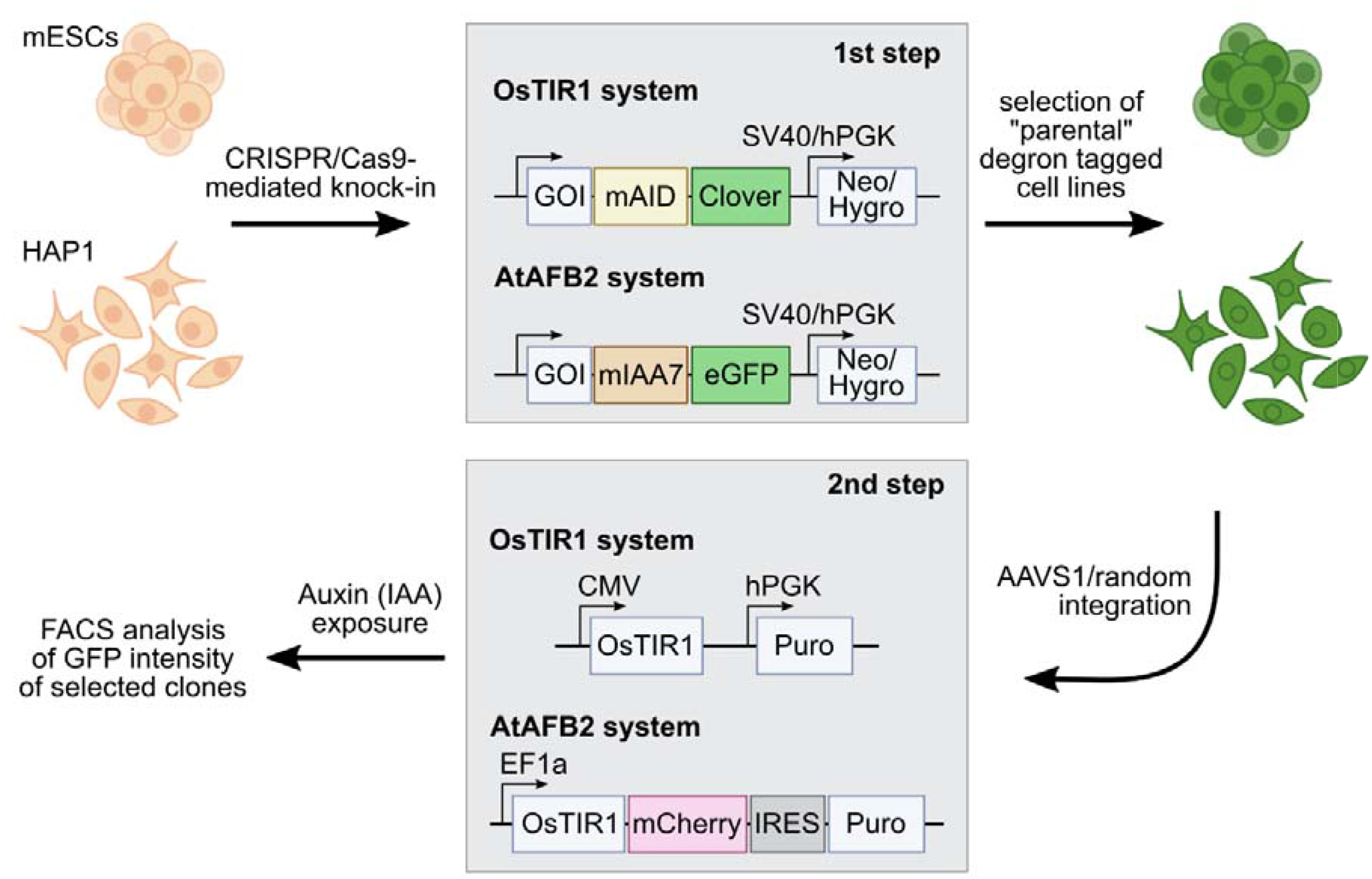
Outline of the AID experiments performed with two AID systems: OsTIR1 and AtAFB2. First step includes targeted integration of the AID degron tags and Neo/Hygro^R^ selection cassette. At second step, selected degron-tagged clones transformed with AFB expression plasmids with Puro^R^ selection cassette. GOI - gene of interest.

In mESCs, the Rad21 gene targeting resulted in 33% homozygous clones; the SMC2 gene - 40%; the CapH gene - 25%; the CapH2 gene - 26%. In HAP1 cells, the Rad21 gene targeting resulted in 11% homozygous clones; the SMC2 gene - 77% (Table 1). We also performed second round modification of one of the mESC Rad21-miniIA 7-eGFP clones to introduce SMC2-miniIAA7-eGFP for double protein degradation. In this case, we could not use Neo/Hygro selection; instead, we visually inspected clones and picked only those with eGFP spread across both nucleus and cytoplasm (in Rad21 clones eGFP is found in the nucleus) (electronic supplementary material Fig. S1).

The correct insertion of the AID cassette was confirmed by long-distance PCR (electronic supplementary material Fig. S2) and Sanger sequencing. Molecular weight of fusion proteins was verified by immunoblotting (in case of Rad21 and SMC2). All analyzed clones were positive for Clover/eGFP expression and had correct protein distribution pattern inside cells, i.e. Rad21-miniIAA7-eGFP and CapH2-miniIAA7-eGFP were confined to the nucleus, while SMC2-miniIAA7-eGFP and CapH-miniIAA7-eGFP resided predominantly in cytoplasm (electronic supplementary material Fig. S1). At least 5 homozygous clones were expanded and frozen for each gene modification and a few (2-3) clones were selected for subsequent transfection with AFB expression vectors.

As far as we know, this is the first application of AID system in human HAP1 cells. This cell line is a powerful tool for genome engineering, because most of the genes are present only in one copy, which simplifies gene modifications. The efficiency of gene targeting was comparable to that of mES cells (Table 1), but required additional selection step, because HAP1 cells have a tendency to spontaneously transit to a diploid state during proliferation [13]. Importantly, ploidy shift interferes with targeting experiments and complicates obtaining homozygous clones after AID cassette targeting. Haploid cells could be selected in two ways. First of all, mixed population of haploid/diploid cells could be reliably sorted by cell size using FACS [13] (Fig. 2A; see also Fig. 2B, where two subpopulation are highlighted with color scheme). Alternately, haploid colonies could be found and picked after simple visual inspection of a plate after antibiotic selection step. Figure 2C demonstrates fluorescent Rad21-miniIA 7-eGFP HAP1 colonies of different ploidy from one of the experiments. HAP1 diploidization at the late passages (after AFB transfections) does not seem to affect AID degradation efficiency.

**Figure 2.**
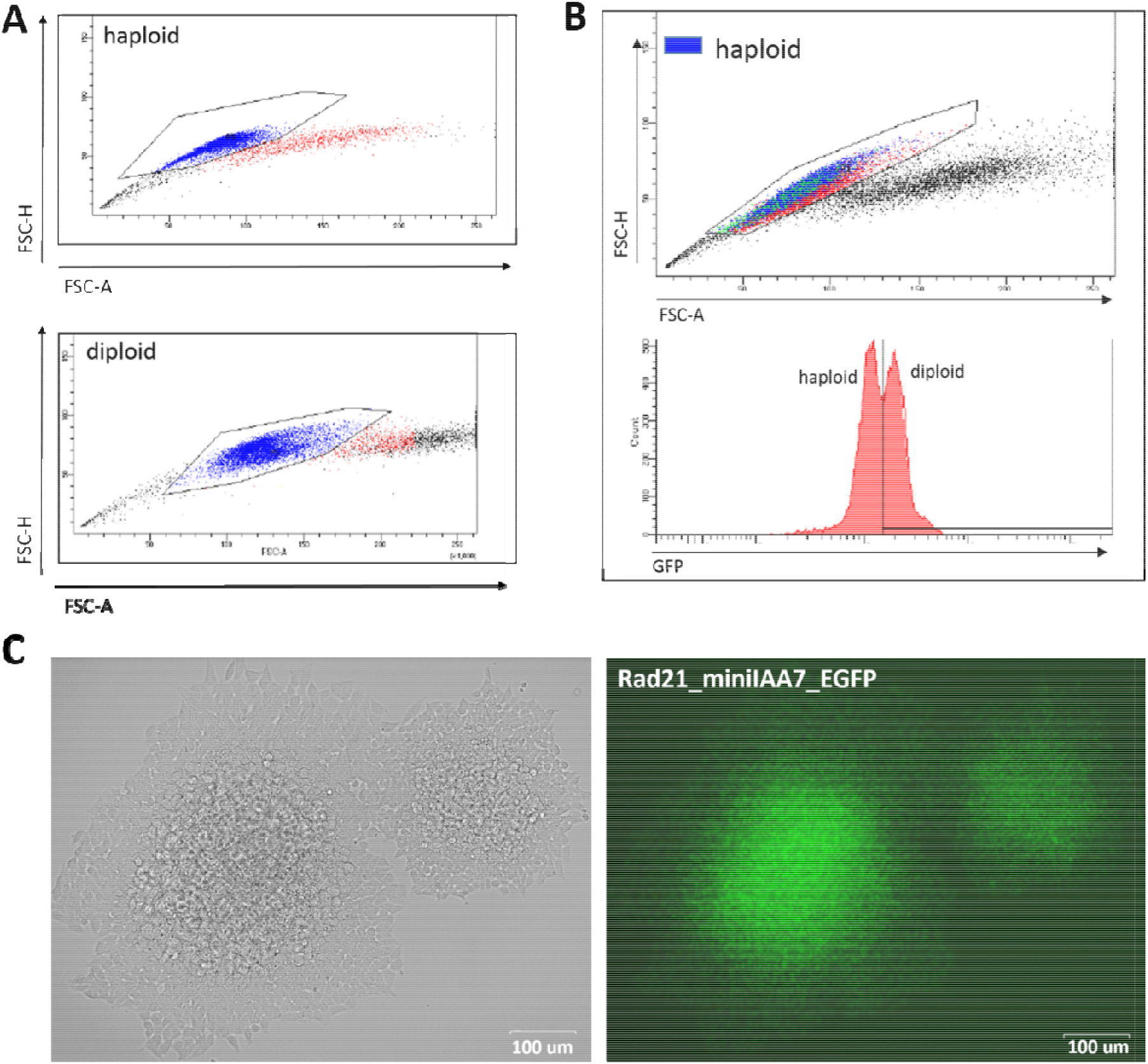
Haploid and diploid HAP1 clones could be reliably separated by cell size using FACS or visual clues. (A) FACS plots for haploid and diploid HAP1 clones. (B) Mosaic HAP1 clone consisting of two subpopulations in equal proportions. Blue/red highlighting indicates haploid/diploid subpopulations, respectively. (C) Two eGFP-positive Rad21-miniIAA7-eGFP clones that show significant cell size differences, making it easy to pick correct clone (haploid clone is on the right).

### Evaluation of the POI degradation efficiency of two AFBs

Selected single homozygous tagged clones (Table 1) were transfected with either CMV-OsTIR1-hPGK-Puro^R^ construct [3], or EF1-AtAFB2-Cherry-IRES-Puro^R^ construct [10], and cultivated in the presence of puromycin for 1 week. Surviving clones with randomly integrated AFBs were expanded and tested for the POI degradation efficiency on FACS. Plot diagrams show distribution of Clover/eGFP fluorescence in mESC and HAP1 clones after 1,5-3 hours of auxin administration (overnight for OsTIR1-mediated degradation) (Fig. 3A,B). OsTIR1 approach using CMV-OsTIR1-hPGK-Puro^R^ was mostly inefficient and required more time to degrade POI without any actual significant depletion. At the same time, AtAFB2 transgene, in which AFB is co-expressed with Puro^R^ and Cherry genes from a single promoter allowed to achieve efficient and rapid degradation. Western blot analysis confirmed FACS results for selected clones (electronic supplementary material Fig. S3).

Clones with the highest level of protein degradation were selected for further studying the dynamics of this process (data for POI = Rad21 and SMC2). We also included HAP1 clones (AtAFB2 system) with intermediate and low sensitivity to auxin to investigate whether longer auxin exposure can enhance protein depletion efficiency.

**Figure 3.**
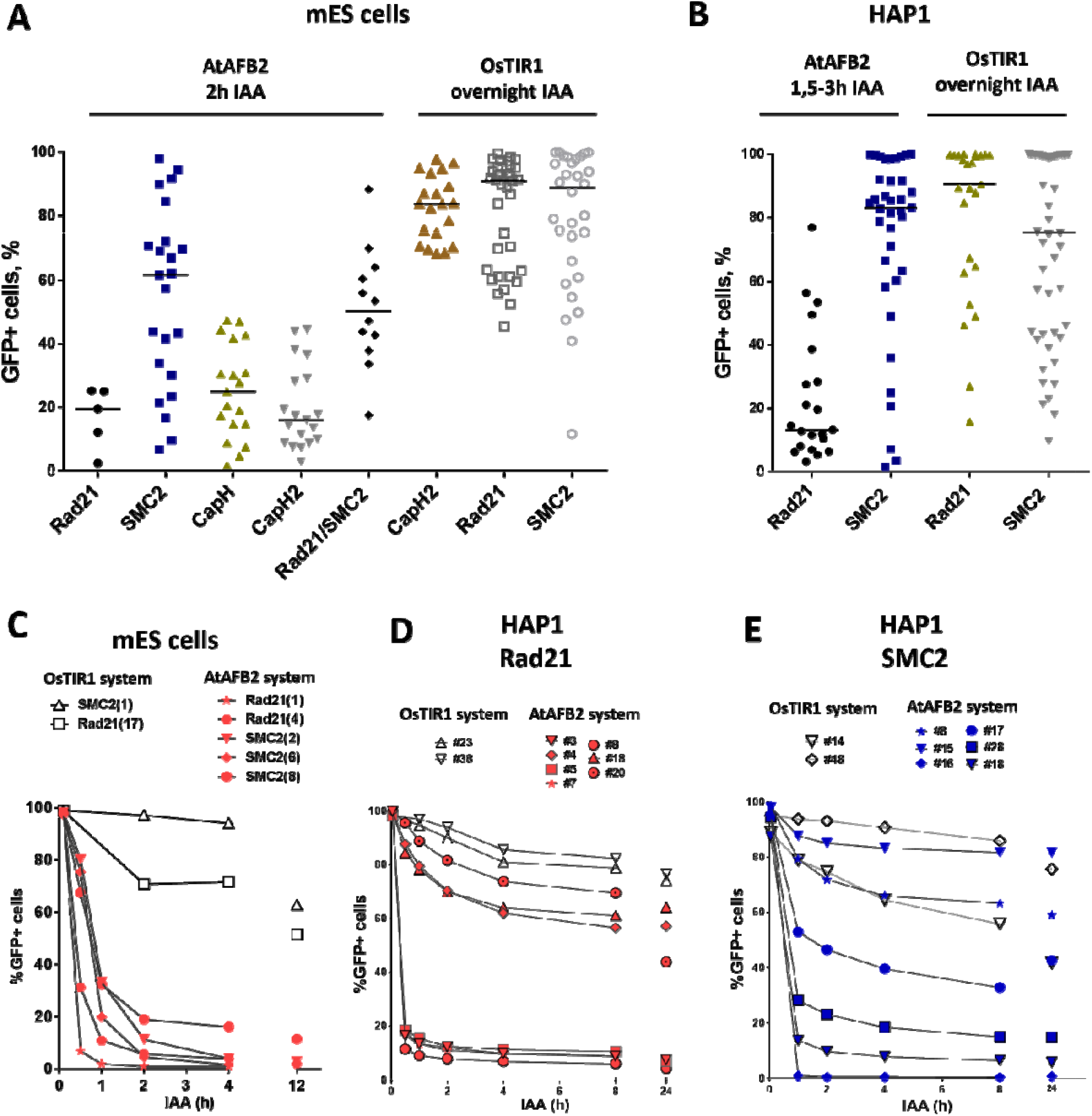
Degradation efficiency and dynamics of POI depletion in mESC and HAP1 clones with randomly integrated OsTIR1 and AtAFB2 constructs. (A) POI degradation efficiency in mESC clones. (B) POI degradation efficiency for HAP1 clones. Black horizontal line shows the median of the group. (C) POI degradation dynamics for selected mESCs clones (Rad21-mAID-Clover, SMC2-mAID-Clover, Rad21-miniIAA7-eGFP, SMC2-miniIAA7-eGFP) with OsTIR1 and AtAFB2. (D) POI degradation dynamics for HAP1 clones (Rad21-miniIAA7-eGFP, Rad21-mAID-Clover) with AtAFB2 and OsTIR1. (E) POI degradation dynamics for HAP1 clones (SMC2-miniIAA7-eGFP, SMC2-mAID-Clover) with AtAFB2 and OsTIR1. Numbers in brackets are individual clones’ designations.

Overall, AtAFB2 cell lines displayed more rapid and efficient depletion of the degron-fused proteins (Fig. 3C-E). For instance, Rad21 levels significantly diminished after 30 minutes of auxin exposure (Fig. 3C, D), which led to collapse of nuclear architecture and cell death after 24 hours in accordance with earlier results. Although AtAFB2-Cherry protein was mostly localized in cytoplasm due to weak NLS, it was enough to rapidly degrade nuclear chromatin factors [10]. Moreover, extended time of auxin exposure does not seem to improve protein degradation efficiency: the most pronounced changes in proteins levels occurred during the first 30-120 minutes and then reached a plateau (Fig. 3E).

AtAFB2 construct contains in-frame Cherry fluorescent marker which is extremely useful to select suitable clones, since Cherry levels reflect AfAFB2 expression. On FACS, we noted strict correlation between Cherry expression and POI degradation efficiency for eGFP signal (data not shown).

### Silencing of the AFB transgene

Although our initial experiment with AtAFB2 AID system demonstrated satisfactory results (Fig. 3), we observed that a subset of cell population remained refractory to auxin and its proportion increased with passages in culture (data not shown). Apparently, decline of POI degradation activity stems from the silencing of AFB constructs. Rad21-miniIAA7-eGFP mES cells and SMC2-miniIAA7-eGFP HAP1 cells bearing randomly integrated AtAFB2 constructs were seeded with low density to generate single-cell founder colonies. After overnight treatment with auxin, we observed both eGFP-positive and negative colonies (Fig.4B). Furthermore, we detected significant number of eGFP/Cherry mosaic colonies (Fig. 4A-B), even though all colonies were derived from single progenitors. Generally, cells with silenced AtAFB2-Cherry construct retained eGFP fluorescence (Fig. 4A-B). For example in Fig. 4A, Cherry^pos^ part of the mESC colony successfully degraded Rad21-miniIAA7-eGFP and started to die, while remaining cells kept expressing Rad21-miniIAA7-eGFP in the absence of AtAFB2-Cherry.

**Figure 4.**
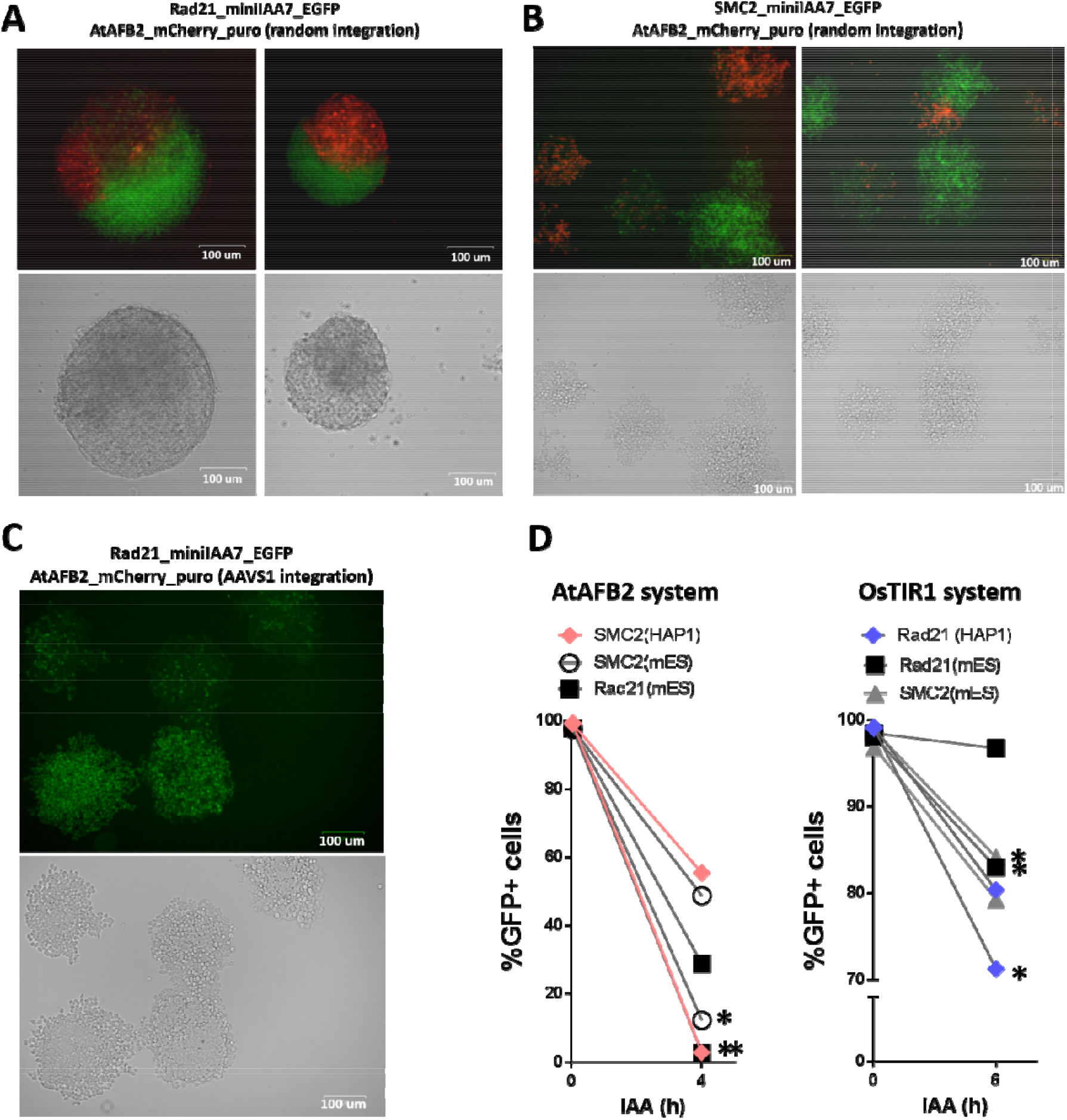
Mosaic expression of AFBs in mESCs and HAP1 cells. One-cell derived colonies of mES cells (A) and HAP1 cells (B) bearing AtAFB2 expression cassette at random genomic loci after overnight auxin exposure. (C) One-cell derived colonies of HAP1 cells with AtAFB2 cassette integration into AAVS1 locus after overnight auxin exposure. (D) Addition of puromycin to culture medium (*) restores the protein degradation efficiency in mosaic clones.

It is well known, that random integration of transgene leads to variable expression due to position effects and silencing. Integration of the OsTIR1 and AtAFB2 transgenes in AAVS1 safe-harbor locus in human cells is considered a sufficient measure to provide stable transgene expression [3,10,12]. We inserted AtAFB2 transgene into AAVS1 locus in one of the HAP1 line with Rad21-miniIAA7-eGFP modification. The AtAFB2 construct has in-frame Cherry selection marker, thus we were able to analyze AtAFB2 expression levels using FACS. Interestingly, all selected Cherry^pos^ clones with correct AtAFB2 insertion were mostly mosaic in expression with 16% Cherry^pos^ cells at best (electronic supplementary material Table S1). None of the 6 clones showed efficient POI depletion in further tests (data not shown). We repeated the experiment with one-cell founder colonies for one of this clones and observed the same outcome with mosaic colonies (Fig. 4C).

Thus, in our experiments, targeted integration of the AFB into AAVS1 safe-harbor locus did not yield any significant advantages over random integration of AFBs. In addition, cells were heterogeneous in AFB expression and ability to degrade AID-tagged protein after auxin exposure.

Our results show that some cells fail to facilitate degradation of target proteins due to the low expression of AtAFB2 protein. Therefore, it is important to keep cells under constant puromycin selection during multiple passages in culture or re-freezing. To demonstrate this, we cultured mES cells bearing Rad21 and SMC2 modifications (AtAFB2 and OsTIR1 system) and HAP1 cells with Rad21 modifications for OsTIR1 system and SMC2 modifications for AtAFB2 system in the presence of 2 µg/ml puromycin for several passages and then analyzed target protein degradation dynamics after auxin treatment (Fig. 4D). As expected, the efficiency of protein degradation was extremely improved in puromycin-treated cells but only for AtAFB2 system (Fig. 4D).

### Culture medium pH levels affect AtAFB2-induced POI degradation

We also inspected whether pH level changes can affect POI degradation efficiencies. Acidification of culture medium pH (6.8 and below) is typically observed in prolonged incubations or in culture mediums with low buffering capacities [14]. We were gradually changing pH values of medium used for cultivating HAP1 Rad21/SMC2-miniIAA7-eGFP clones and later detected eGFP fluorescence on FACS, similar to previous experiments. Normal and mildly acidic pH (6.3-7.4) did not affect POI degradation efficiency, while higher pH values (8.6-8.8) effectively blocked POI degradation (Fig. 5A,B). Auxin analog NAA (1-naphthaleneacetic acid) showed similar pH sensitivity (Fig. 5B). Therefore, physiological pH changes in cell culture medium do not affect auxin-dependent protein degradation efficiency.

**Figure 5.**
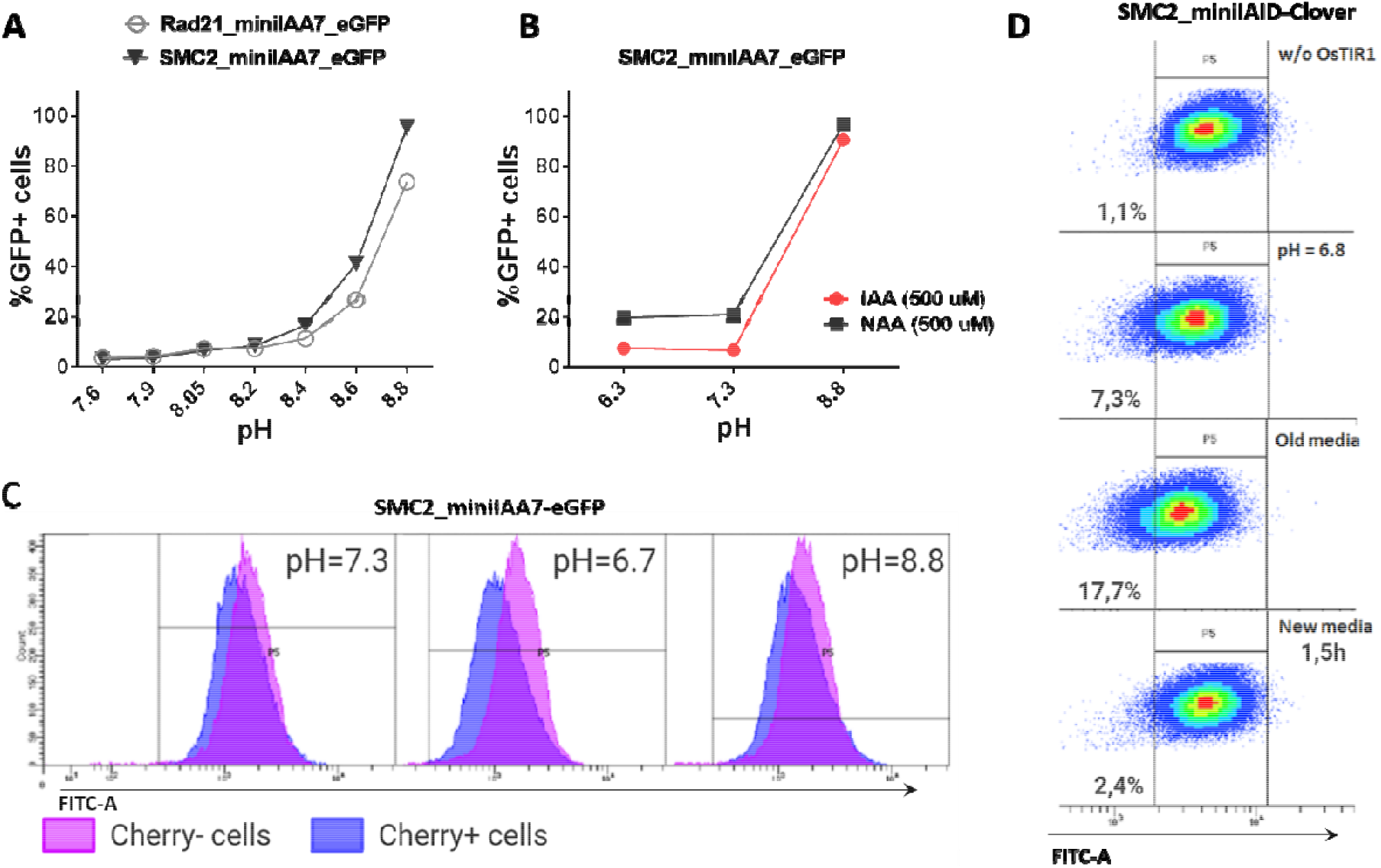
Effects of pH on AtAFB2 and OsTIR1 AID degradation efficiency. (A) High pH levels inhibit auxin-induced eGFP degradation in HAP1 clones with Rad21 and SMC2 modifications. (B) IAA and NAA auxin analogs show similar sensitivity to pH changes (HAP1 clones). (C) eGFP levels in mosaic HAP1 clone SMC2-miniIAA7-eGFP. Blue color indicates eGFP levels in Cherry^pos^ subpopulation; purple color - Cherry^neg^ subpopulation. The shift between two peaks illustrates minor basal degradation induced by AtAFB2 in Cherry^pos^ cells. (D) The shift of mES cell population (OsTIR1 system) towards left on FITC-A axis in acidic media.

Nevertheless, low pH might influence basal degradation in AtAFB2 AID system. We inspected mosaic HAP1 clone with miniIAA7-tagged SMC2, which consists of mixed Cherry^pos^/Cherry^neg^ subpopulations, caused by the silencing of the AtAFB2 expression in some cells (Fig. 4B). Generally, Cherry^pos^ cells has slightly weaker eGFP fluorescence profile (blue, (Fig. 5C)) with a shift between two peaks, probably reflecting minor basal degradation. At normal pH or high pH (no AtAFB2 activity), eGFP level shifts remain similar. However, changing pH value to 6.7 led to a slight decrease in eGFP fluorescence in Cherry^pos^ cells, but not in Cherry^neg^ cells (Fig. 5C, middle), which might hint at increased basal degradation at low pH. We observed the same slight shift for OsTIR1 AID system while culturing cells in acidic pH medium, including culture media in which cells were growing overnight (old media) (5D).

## DISCUSSION

The auxin-inducible degron (AID) system consists of two molecular components. To begin with, a gene of interest has to be tagged with AID degron domain and selection cassette with one of two markers for double selection (e.g. NeomycinR/HygromycinR) [3]. Obtaining homozygous clones using double selection is convenient and highly efficient. In our experiments, for all six loci tagged (hRad21, hSMC2, mRad21, mSMC2, mCAPH, mCAPH2) overall efficiency ranged from 11 to 70% (Table 1). The only inconvenience that we faced was spontaneous diploidization of HAP1 cells (Fig. 2). To avoid problems with obtaining homozygous integrations, it is important to select haploid clones by visual inspection of colonies or by FACS [13]. Additionally, one should check copy number of the target loci in advance using real-time PCR or analogous methods, because chromosome number may vary between HAP1 clones [15].

The key step to successful AID experiments is high and stable AFB (receptor F-box protein i.e OsTIR1 and AtAFB2) expression. Our pilot experiments with mAID/OsTIR1 system involved six proteins of interest (POIs) (Fig. 1) and followed a popular protocol [3]. Using this approach, we obtained many clones with correct gene modifications (Table 1), but never succeeded in getting reliable degradation with CMV-OsTIR1-Puro expression cassette (Fig. 3A-B). Most experiments on cultured cells used random OsTIR1 integration [4,5,16–18]. In theory, integration of the AFBs in the safe-harbor locus should provide sufficient expression levels. This approach is recommended in majority of protocols where many suitable loci (H11, TIGRE, AAVS1, ROSA26) were extensively characterized [4,7,19]. The downsides of the safe-harbor targeting approach are extra experimental manipulations, restricted expression levels, and susceptibility to silencing, just as in randomly integrated transgene constructs (Fig. 4). For instance, TIGRE safe-harbor locus expression may vary after differentiation [4], which is critical for ES cell experiments. During our experiments, we tested both of these approaches, and found no significant difference between targeted and random integration. Therefore, generating a set of clones with highly expressed randomly integrated AFB transgenes might be more convenient, while providing clones with diverse expression levels suitable for any specific POI.

Surprisingly, we found that selected clones with efficient POI degradation became refractory to auxin after some time in culture due to AFB silencing (Fig. 4). For example, we observed mosaic mESC colony where roughly half of the cells lost Cherry expression (low AtAFB2) and couldn’t degrade Rad21-miniIAA7-eGFP (Fig. 4A-B). Mosaic silencing happens almost inevitably in most clones, which is unacceptable for depletion endeavors requiring 0-10% POI levels. Continuous supplementation of puromycin to culture medium prevents AFB silencing in further experiments (Fig. 4D). We feel that this fact is either omitted or not stressed enough in most published protocols [3,9,11,12]. Our observations also warn against removing selection cassette after AFB integration in safe-harbor loci [7]. In future, exclusion of the AFB expression step could help to improve the AID experience [20], but the method is still under development.

Comparing the two AID systems directly in our case could be difficult, because AFB expression vectors differ between protocols [3,10]. AtAFB2 has in-frame fluorescent Cherry reporter useful for selection of high AFB expressing clones during visual colony examination. However, it doesn’t prevent silencing of the Cherry-expressing cells later (Fig. 4). In AtAFB2 construct, Puro^R^ gene is linked to AFB via IRES, while in CMV-OsTIR1-Puro vector Puro^R^ expression is driven from independent promoter (Fig. 1). The latter could hinder puromycin selection. Nevertheless, after AFB transfection and 1-week puromycin selection AtAFB2 clearly wins over OsTIR1 in terms of POI degradation efficiency (Fig. 3A-B). Other factors, such as absence of basal depletion and degradation dynamics were also in favor of AtAFB2 system (Fig. 3C-E; Fig. 5C). Recent study reports that basal degradation in OsTIR1 AID could be countered by overexpressing ARF protein (Sathyan et al., 2020). Unfortunately, this approach is not applicable for the popular miniAID (mAID) degron system, because mAID lacks ARF-interacting domains [12].

It is known, that the binding affinity of auxin with its receptors would be pH-dependent [21]. To our knowledge, how pH levels affect auxin-dependent degradation in cultured mammalian cells was never reported before. We showed in HAP1 cell lines that increasing pH from 7.4 to 8.8 completely blocks AtAFB2-dependent degradation, while decrease of pH doesn’t affect AtAFB2 (Fig. 5). Such pH-based switch could be a simple lever to temporarily block POI degradation during experiments. AID pH sensitivity might be worth testing in the context of dual degradation approach like in alternative coronatine-dependent JAZ degron/COI system combined with OsTIR1 [22,23].

## CONCLUSION

In conclusion, we would like to outline AID approach that showed success in our hands. Our experience is based on evaluation of the OsTIR1 and AtAFB2 systems in mES and human HAP1 cells and might help readers to avoid common pitfalls. One should aim to generate bi-allelic tag modifications first using CRISRP/Cas9 and miniIAA7 donor vectors with double selection markers. Then select a few clones validated with long-distance PCR and Sanger sequencing, and randomly insert AtAFB2 cassette with in-frame Puro^R^ and Cherry selection markers. Usually 10-20% of puromycin-resistant AtAFB2 clones display high POI degradation efficiency. Always use puromycin selection during experiments to avoid AtAFB2 silencing.

## MATERIAL AND METHODS

### Cell culture

All cell lines were grown at 37°C with 5% CO_2_ and passaged every 2–3 days. Hap1 cells (Horizon Discovery) were cultured in IMDM (ThermoFisher) containing 10% FBS (Gibco), 1mM L-glutamine (Sigma), 0.5 mM NEAA (Gibco), penicillin/streptomycin (100 U ml^-1^ each). Mouse ES cells (DGES1 line) established in our laboratory [24]) were cultured on a gelatin surface under 2i condition (1 μM PD, 3 μM CHIR) in DMEM (ThermoFisher), supplemented with 7.5% ES FBS (Gibco), 7.5% KSR (Gibco), 1 mM L-glutamine (Sigma), NEAA (Gibco), 0.1 mM β□mercaptoethanol and LIF (1000 U/ml, Polygen), penicillin/streptomycin (100 U ml^-1^ each). 3-Indoleacetic acid (IAA, auxin) (Merck, I2886) and 1-Naphthaleneacetic acid (NAA) (Merck, N0640) were dissolved in NaOH (1M) solution to concentration of 500 mM, aliquoted, stored at −20°C, and used immediately after thawing. For all experiments, cells were incubated with 500 μM indole-3-acetic acid (IAA) (Sigma) for the indicated times.

For testing the pH-responsive auxin-inducible degradation of POI, cell culture media with a pH range of 6.3-8.8 was prepared and used immediately. The cells were harvested no longer than 30 min post auxin exposure, because alkaline pH rapidly returns to neutral values in CO2-rich atmosphere.

### Construction of the AID donor vectors and site-specific degron integration

Donor vectors with homology arms, degron tag (mAID or miniIAA7), Clover/eGFP, and selection cassette (either Neo^R^ or Hygro^R^) were assembled in one reaction using NEBuilder HiFi kit (NEB) in pMK289 backbone [3]. Homology arms for hRad21 (Gene ID: 5885), hSMC2 (Gene ID: 10592), mRad21 (Gene ID: 19357), mSMC2 (Gene ID: 14211), mCapH (Gene ID: 215387), mCapH2 (Gene ID: 52683) genes were PCR amplified from human and mouse genomic DNA with Q5 polymerase (NEB). The lengths of homology arms are featured in electronic supplementary material Table S2. Other functional elements were amplified from plasmids pMK289 (mAID-mClover-NeoR) (Addgene #72827), pMK290 (mAID-mClover-HygroR) (Addgene #72828) and pSH-EFIRES-B-Seipin-miniIAA7-mEGFP (Addgene #129719) [3,10]. For auxin receptor F-box proteins (AFBs) overexpression, OsTIR1 and AtAFB2 plasmids were used (AtAFB2) (pMK232 CMV-OsTIR1-PURO, Addgene #72834; pSH-EFIRES-P-AtAFB2-mCherry-weakNLS, Addgene #129717) [3,10]. For gene tagging and AAVS1 locus modification, cells were transfected with donor vectors, Cas9 (Addgene #41815) and corresponding sgRNA-expressing plasmids (Addgene #41824, backbone). Sequences of sgRNAs are provided in electronic supplementary material Table S2.

### Generation of transgenic cell lines with auxin-dependent degradation of POI

HAP1 cells were electroporated at parameters of 1200 V, 30 ms, 1 pulse using Neon Transfection system (Invitrogen), according to the manufacturer’s instructions. The day after electroporation cell were seeded in 100-mm dishes with low density and expanded for 24 hours, followed by selection with 500 μg/ml G418 (for Rad21 targeting), or 800 μg/ml Hygromycin B (for SMC2 targeting). Mouse ES cells were transfected by Lipofectamine 3000 (Invitrogen) in OPTI-MEM (ThermoFisher), following supplier recommendations. 24 hours post-transfection cells were seeded on 60-mm dishes in culture medium containing 300 μg/ml G418 and 200 μg/ml Hygromycin B for selection of homozygous tagged clones [3]. After 10-14 days of selection, resistant colonies of HAP1 and mouse ESCs expressing green fluorescent protein (eGFP) were manually picked for subcloning and further analysis. Most of the surviving clones were confirmed to harbor the correct insertion of degron tags by genomic PCR, Sanger sequencing and western blotting. In, addition, the resulting HAP1 clones were stained with propidium iodide (PI) and their DNA ploidy was determined by FACS [13]. Several clones for each target gene were chosen for integration of AtAFB2 or OsTIR1 expressing vectors and transformed using identical settings described for the previous step. Selection of the AFB positive clones was performed using 1 μg/ml puromycin for 1 week. Finally, several mESCs and HAP1 clones with homozygous degron tags and AFBs insertions were tested for efficiency of auxin-induced protein depletion, using microscopy, FACS and Western blotting.

### PCR genotyping

Cells were lysed in PBND lysis buffer (10 mM Tris-HCl, 50 mM KCl, 2.5 mM MgCl2, 0.1 mg/ml gelatin, 0.45% NP-40, and 0.45% Tween20, pH 8.3) containing proteinase K (200 μg /ml) for 1h at 55°C followed by Proteinase K inactivation for 10 minutes at 95°C. The target regions were amplified by PCR with either Taq-HP DNA Polymerase (Biospecifika Novosibirsk, Russia) or LongAmp Taq DNA Polymerase (NEB) (primers sequences could be found in electronic supplementary material Table S3). The parameters were as follows: 95°C for 30 s, then 35 cycles of 94°C for 10 s, 60°C for 30 s, 72°C for 1 min (1 min/kb) and a final step at 72 °C for 5 min for Taq-HP DNA Polymerase; and 94°C for 30 s, then 30 cycles of 94°C for 10 s, 60°C for 30 s, 65°C for 3-6 min (1 min/kb) and a final step at 65 °C for 5 min, for LongAmp Taq DNA Polymerase. The amplified products were analyzed by agarose gel electrophoresis.

### Western blotting

Cells were washed twice with PBS and scrapped from the surface in presence of RIPA buffer (50 mM Tris-HCl pH 8, 150 mM NaCl, 1% Triton X100, 0.5% sodium deoxycholate, and 0.1% SDS) containing the protease inhibitor cocktail (1x Complete ULTRA (Roche), 1x PhosSTOP (Roche), 5 mM NaF (Sigma)). After that, cells were sonicated by three 10 s pulses at 33-35% power settings. Lysates were centrifuged at 18000 g for 20 minutes at 2°C, frozen and stored at −80°C. The protein concentrations in cell lysates were quantified using Pierce BCA Protein Assay Kit (ThermoFisher). Equal amounts (25 μg) of total protein were separated on 10% SDS-PAGE, and then transferred onto Immun-Blot PVDF membrane (Bio-Rad). After blocking in 5% milk/TBST for 2 h, membrane was incubated with primary antibodies against RAD21 and SMC2 (#12673/#8720 Cell Signaling technology) at 4°C overnight. At the following day, membranes were incubated with horseradish peroxidase-conjugated secondary antibodies (#7074 Cell Signaling technology) for 2 hours at 25°C. Detection was performed with SuperSignal West Pico PLUS Chemiluminescent Substrate (#34580, ThermoFisher) and Chemidoc XRS Imaging system (Bio-Rad).

GraphPad Prism 6.0 was used to design graphs and analyze quantitative data. The results are presented as mean values ± standard deviations.

## Data accessibility

This article has no additional data.

## Authors’ contributions

NB conceived the study. AY, AS, SA and TS collected and analyzed the data. AY and AS wrote the manuscript. All authors read and approved the final manuscript.

## Acknowledgements

The authors thank the Collective Center of ICG SB RAS “Flow Cytometry” for the access to research infrastructure. We also express our gratitude to Dr. A.G. Menzorov and Collective Center of ICG SB RAS “Collection of Pluripotent Human and Mammalian Cell Cultures for Biological and Biomedical Research” for DGES1 cell line.

---

**Supplementary Figure S1.**
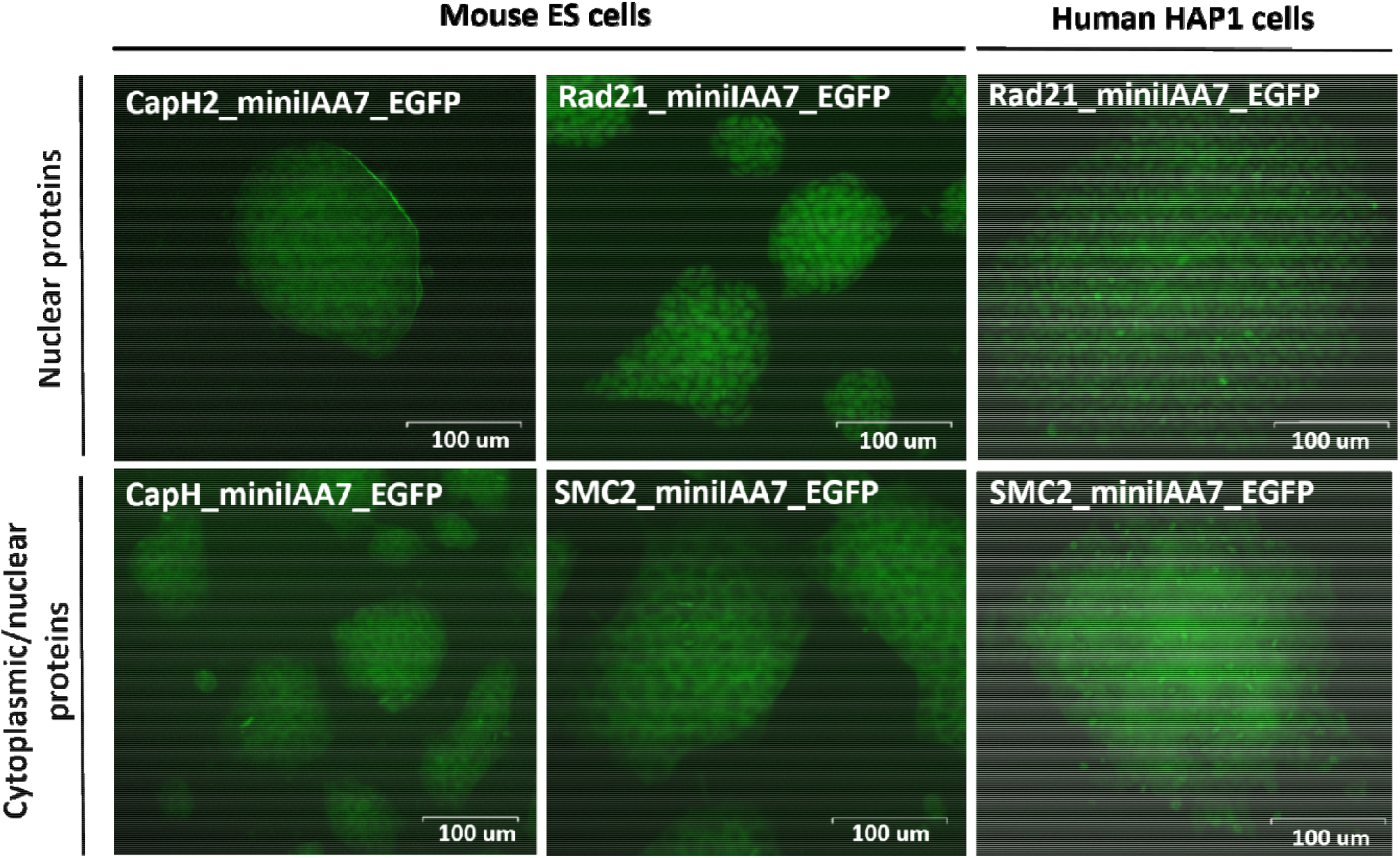
Distribution of the POI-miniIAA7-eGFP protein in homozygous tagged mESCs lines and HAP1 lines.Top pictures: Nuclear localization (Rad21 and CapH2 clones). Bottom pictures: Predominantly cytoplasmic localization (SMC2 and CapH clones). Thick bands visible in some of the cells label mitotic spindles, which are covered with condensins during division.

**Supplementary Figure S2.**
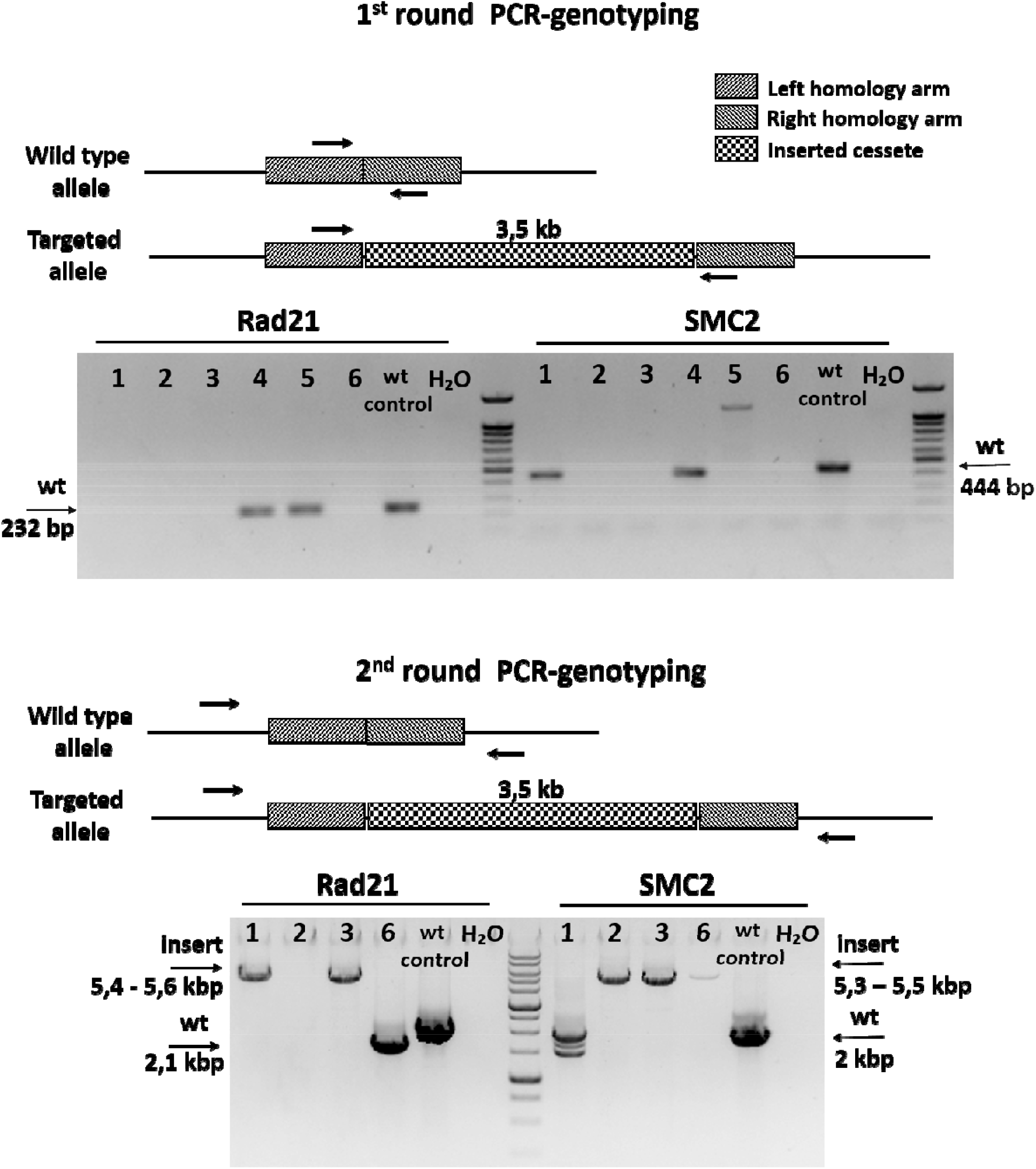
PCR genotyping strategy for selecting correctly AID tagged genes (Rad21 and SMC2 mESC clones; miniIAA7 tag). 1^st^ round: primers complimentary to the modified loci amplify wild-type (wt) alleles (300-500 bp). The lack of amplicons indicates either targeted cassette insertion or the large deletions involving the primer-binding sites. 2^nd^ round: primers beyond the donor plasmid homology arms were used in long-distance PCR (5-6 kbp). Indicated amplicon sizes confirm the correct integration of constructs. Amplicons for wt and targeted alleles (AID cassette insertion) are marked with arrows. Apparently, some wt alleles carry deletions at the modified loci (Rad21 N2,6; SMC2 N1).

**Supplementary Figure S3.**
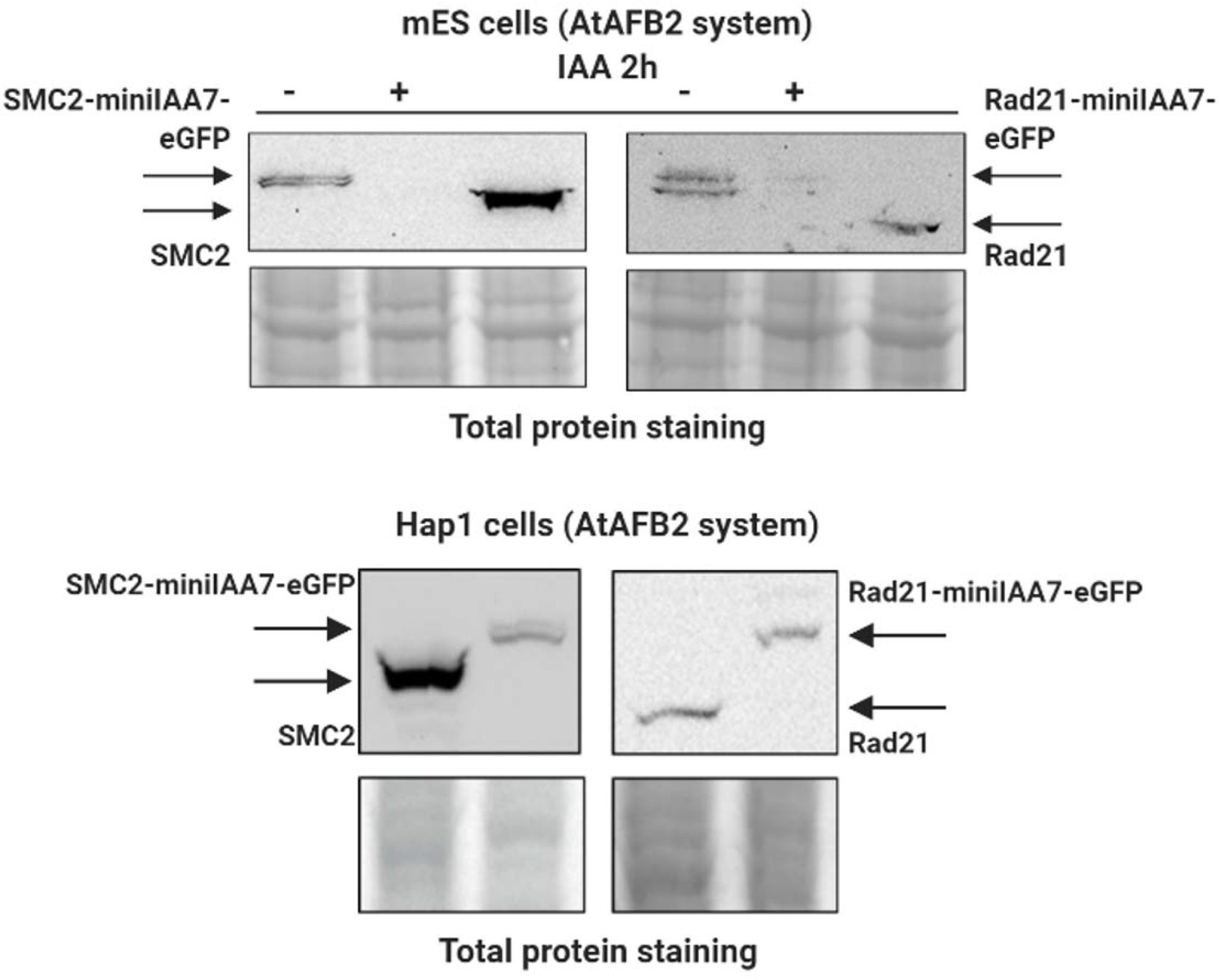
Western blot analysis for SMC2 and Rad21 proteins before and after insertion of degron tags in mESCs and HAP1 cell clones. (+)-degradation of POI-miniIAA7-eGFP after addition of auxin (IAA). (−)-no auxin.

**Supplementary Table S1.** Integration of the AtAFB2 in the AAVS1 locus results in mosaic clones (%, Cherry) with low degradation efficiency.

| **Clone number** | **Ploidy** | **%, Cherry+ cells** | **AAVS1 integration** |
| --- | --- | --- | --- |
| 20 | hap | 16,2 | + |
| 34 | hap | 12,5 | + |
| 54 | hap | 12,4 | + |
| 58 | hap | 6,3 | + |
| 64 | hap | 6 | + |
| 72 | hap | 1,9 | + |

**Supplementary Table S2.**
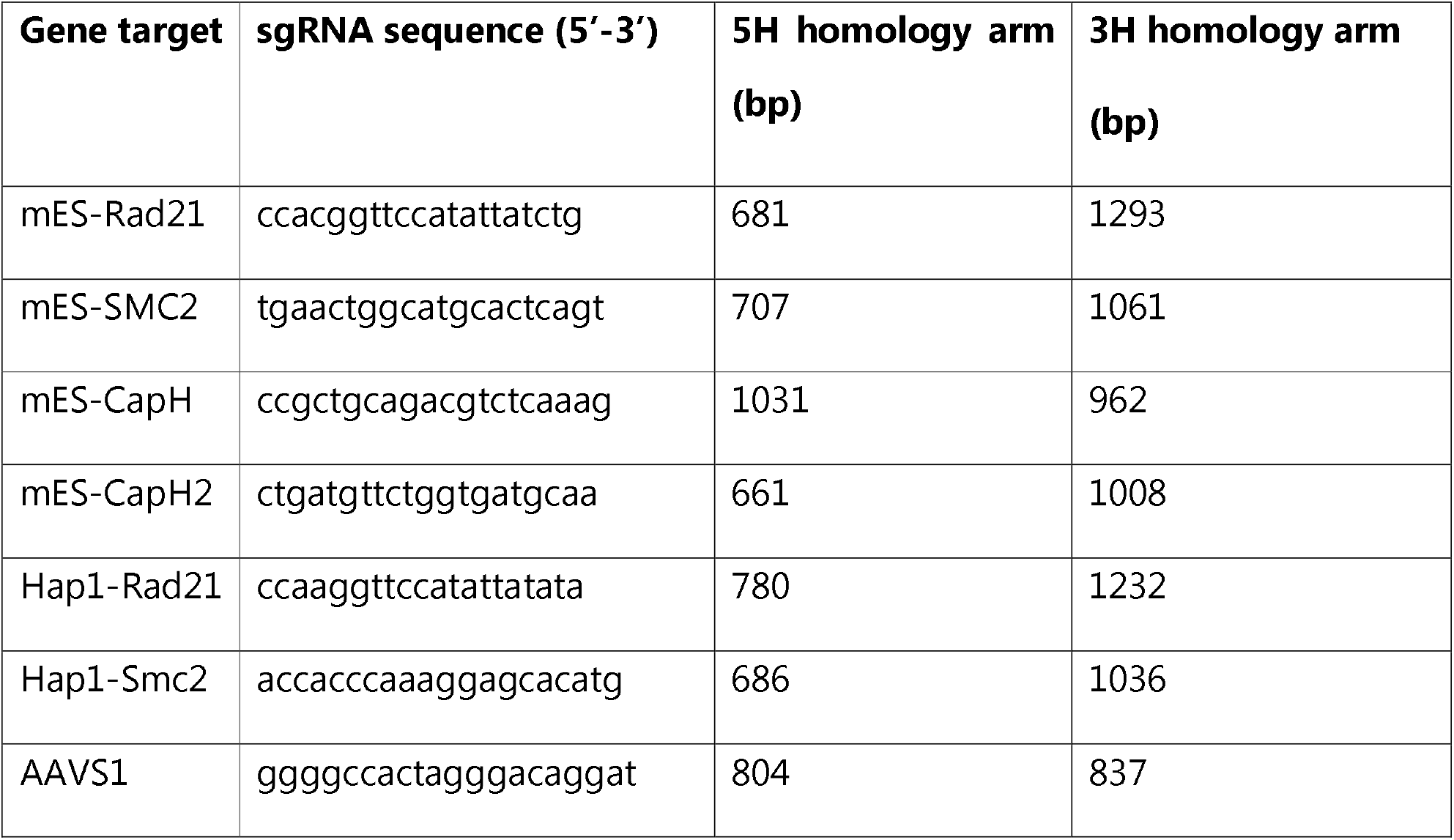
Sequences of sgRNAs and donor vector arms lengths used in AID targeting experiments.

**Supplementary Table S3.**
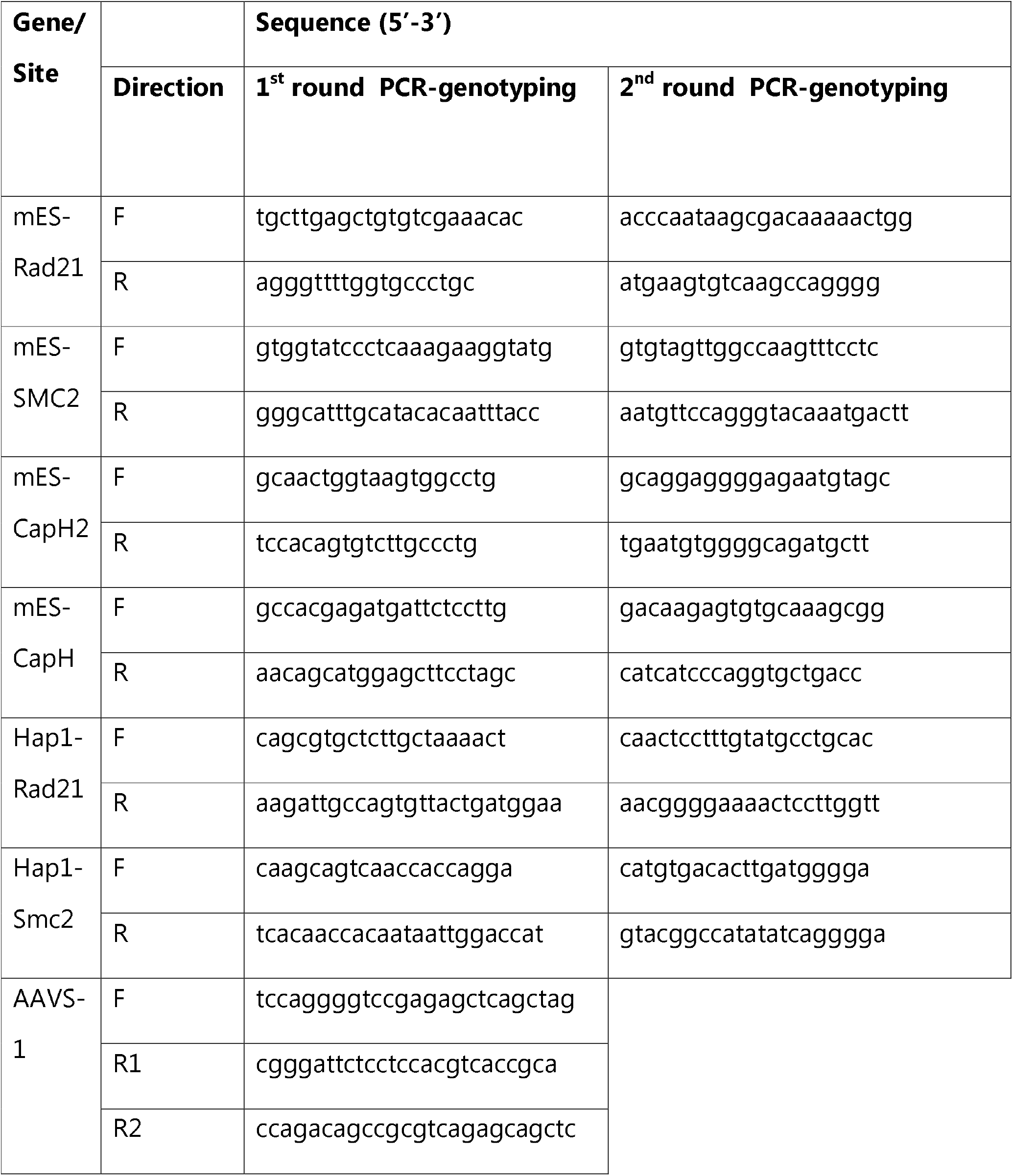
Oligonucleotides used for genotyping of modified clones.

